# Vault RNAs aid viral infection by facilitating nuclear export of hnRNP C and ELAVL1

**DOI:** 10.1101/2024.10.07.616963

**Authors:** Jorn E. Stok, Sander B. van der Kooij, Dennis Gravekamp, Laurens R. ter Haar, Jasper W. de Wolf, Clarisse Salgado-Benvindo, Tessa Nelemans, Bart J.M. Grijmans, Rayman T.N. Tjokrodirijo, Arnoud H. de Ru, Peter A. van Veelen, Martijn J. van Hemert, Marjolein Kikkert, Annemarthe G. van der Veen

## Abstract

Vault RNAs (vtRNAs) are a family of four small non-coding RNAs (ncRNAs) that are ubiquitously expressed in many eukaryotes and that regulate multiple cellular pathways. Their expression is increased upon infection with various DNA and RNA viruses. This suggests they are either co-opted by the virus to aid replication or function as an antiviral restriction factor. However, their precise molecular function remains unclear. Here, we show that replication of picornaviruses, alphaviruses, and beta-coronaviruses broadly enhances vtRNA expression. We find that genetic loss of vtRNAs inhibits replication of Sindbis virus (SINV) and encephalomyocarditis virus (EMCV), independent of the antiviral type I interferon (IFN) response. A proteomic screen uncovered the vtRNA interactome and revealed that vtRNAs associate with RNA binding proteins ELAVL1 and hnRNP C in uninfected and infected cells. VtRNAs facilitate the translocation of ELAVL1 and hnRNP C from the nucleus to the cytoplasm in infected cells, an event that is required for efficient viral replication. Moreover, hnRNP C and ELAVL1 fail to associate with viral RNA in the cytosol of SINV-infected cells in the absence of vtRNAs. Together, our findings reveal a novel molecular mechanism by which vtRNAs exert proviral activity during the course of SINV and EMCV infection, which opens up new avenues for therapeutic targeting to fight infectious diseases.

## Introduction

Despite the small size of viral genomes, viruses have evolved refined mechanisms to exploit the cellular machinery of the host cell to ensure their own replication and survival, as well as to evade antiviral defense pathways. It is essential to understand such mechanisms as they provide potential targets for novel antiviral medication. Countless examples describe how viral proteins hijack host proteins to manipulate cellular processes^1–4^. However, only a small part of the human genome contains protein-coding genes, while large portions of the ‘non-coding’ genome are also actively transcribed. The resulting non-coding RNAs (ncRNAs) have emerged as crucial players and regulators in diverse cellular pathways^5,6^. Viruses can manipulate ncRNAs and use these to their own advantage to sustain their replication. Conversely, ncRNAs may promote antiviral defense mechanisms to limit viral spread^7,8^.

Vault RNAs (vtRNAs) are a family of small ncRNAs of 89-121 nucleotides in length that are transcribed by RNA polymerase III, ubiquitously expressed in eukaryotic cells, and conserved throughout evolution^9^. In humans, four highly homologous vtRNAs are expressed: vtRNA1-1, vtRNA1-2, vtRNA1-3, and vtRNA2-1. Instead, mice only express a single vtRNA (mvtRNA)^9^. VtRNAs were originally discovered as components of the vault complex, the largest ribonucleoprotein complex in eukaryotic cells^10^. Although the assembly and components of the vault complex are well-documented, its function remains elusive. The human vault complex is composed of multiple copies of three proteins, namely the major vault protein (MVP), vault poly(ADP-ribose) polymerase (vPARP), and telomerase-associated protein 1 (TEP1)^10–12^. The latter binds vtRNAs and docks them to the vault complex^13–15^.

However, only an estimated 5% of the cellular pool of vtRNAs is tethered to the vault complex via TEP1. The remaining 95% exists freely in the cytoplasm and nucleus, which suggests they fulfill biological functions independent of the vault complex^16–18^. Indeed, vtRNAs are involved in various cellular processes^19,20^. For example, vtRNA1-1 negatively regulates the multimerization of the ubiquitin receptor p62 (or sequestosome-1 (SQSTM1)), thereby inhibiting autophagy^21–23^. VtRNA1-1 also confers resistance to apoptosis and to chemotherapeutic treatment, highlighting its pro-survival role^24–27^. Additionally, human vtRNA1-1 and mouse mvtRNA are involved in synapse formation in neurons^28,29^. VtRNA2-1 (also known as nc886) binds to the RNA-binding protein (RBP) ELAVL1 (also known as HuR). It prevents ELAVL1 from stabilizing mRNAs encoding intracellular junction proteins, which in turn impairs cell-cell contact and epithelial barrier function in the intestine^30^. Additionally, vtRNA2-1 may interact with and suppress activation of the double-stranded RNA (dsRNA) receptor protein kinase R (PKR), which controls protein translation and cell growth^31–33^. Unlike vtRNA1-1 and vtRNA2-1, the molecular function and interaction partners of vtRNA1-2 and vtRNA1-3 are largely unknown. Lastly, vtRNAs may be post-transcriptionally modified by incorporation of a 5-methylcytosine (m^5^C) modification. This leads to Dicer-dependent cleavage of these RNAs into small vtRNA fragments, which can subsequently interact with Argonaute and regulate gene expression in a miRNA-like fashion ^34–37^.

The expression of vtRNAs is strongly increased upon infection with a select number of RNA and DNA viruses, including Influenza A virus (IAV), Epstein-Barr virus (EBV), and herpes simplex virus 1 (HSV-1)^18,24,38,39^. Knockdown of vtRNAs impairs replication of IAV as a result of increased PKR activation and NF-κB signaling^39^. In addition, overexpression of vtRNA1-1 in B cells enhances EBV establishment^24^. Finally, vtRNA2-1 enhances adenovirus replication by promoting nuclear entry of the viral genome^40^. These studies suggest that vtRNAs have proviral activity and are hijacked by replicating viruses. It is not clear whether the assumed proviral activity of vtRNAs is broadly applicable to a diverse range of viruses; nor if this activity applies to all four vtRNAs or to a subset. In addition, the precise mechanistic bases underlying the potential proviral capacity of vtRNAs remain largely unknown. Conversely, some studies have indicated that vtRNAs bind the viral RNA sensor retinoic acid-inducible gene I (RIG-I) during infection with Dengue virus or Kaposi’s sarcoma-associated herpesvirus (KSHV)^41,42^. In such instances, vtRNAs may serve as a host-derived ligand that enables RIG-I activation and the downstream antiviral type I interferon (IFN) response, and act as a viral restriction factor^41,42^. Here, we investigated how vtRNAs affect viral infection and through which mechanisms this occurs. We show that viruses of distinct families, i.e. picornaviruses, alphaviruses, and beta-coronaviruses, broadly induce vtRNA expression. Using genetic approaches, we demonstrate that vtRNAs have a proviral effect in alpha- and picornavirus infections that does not depend on modulation of the type I IFN response. Using a proteomics approach, we define the protein interactome of vtRNA1-1, vtRNA1-2, and vtRNA1-3 and find that vtRNAs strongly associate with the RBPs hnRNP C and ELAVL1 in uninfected and infected cells. Finally, we show that the loss of vtRNAs affects the nuclear-cytoplasmic shuttling of hnRNP C and ELAVL1 during viral infection, which is a prerequisite for successful viral replication. Our findings provide novel molecular insight into the function of vtRNAs and their exploitation by distinct viruses.

## Results

### Infection with various RNA viruses increases vtRNA expression

Several studies have reported increased vtRNA expression during infection with certain viruses, such as Epstein-Barr virus (EBV) and Influenza A virus (IAV)^18,24,38,39^. We set out to investigate whether vtRNA expression is also upregulated during other viral infections. We infected the human alveolar carcinoma cell line A549 with encephalomyocarditis virus (EMCV), a cardiovirus from the *Picornaviridae* family, and assessed vtRNA expression 24 hours (h) post-infection by RT-qPCR. We noticed that the expression of commonly used housekeeping genes, such as *ACTB*, *GAPDH,* and *RPS13*, were generally affected during viral infection, including EMCV infection (Fig. S1A). We therefore chose to normalize RT-qPCR data using *RN18S1*, an RNA polymerase I transcript that remains stably expressed throughout the course of viral infection. We found that expression of vtRNA1-1, vtRNA1-2, and vtRNA1-3 increased on average 11- to 14-fold upon EMCV infection, while the expression of vtRNA2-1 remained unchanged (Fig. 1A). Similarly, infection of A549 cells with Mengovirus, a closely related cardiovirus, induced expression of vtRNA1-1, vtRNA1-2, and vtRNA1-3, but not vtRNA2-1, about 10-fold (Fig. 1B). Infection of human embryonic kidney fibroblasts (HEK293) or lung fibroblasts (MRC5) with three different alphaviruses, namely Sindbis virus (SINV), Chikungunya virus (CHIKV), and Venezuelan Equine Encephalitis Virus (VEEV), all enhanced expression of vtRNAs (Fig. 1C-E). Additionally, Middle Eastern respiratory syndrome-related coronavirus (MERS-CoV) strongly induced vtRNA expression in MRC5 cells (Fig. 1F). Note that the overall magnitude of vtRNA upregulation differed per virus and vtRNA. For example, while most viruses did not impact vtRNA2-1 expression, infection with VEEV and MERS-CoV led to an 8-fold and 31-fold increase in vtRNA2-1 expression, respectively (Fig. 1A-F). To test whether murine viruses also upregulate the expression of mvtRNA, we infected the mouse embryonic fibroblast cell line 17cl1 with the murine coronavirus mouse hepatitis virus (MHV). We observed upregulation of mvtRNA 16h post-infection (Fig. 1G). Importantly, not all viral infections caused increased vtRNA expression. Infection of THP-1 macrophages with reovirus, or A549 cells with adenovirus (AdV), Usutu virus (USUV), or West Nile virus (WNV) did not increase vtRNA expression at 24h post-infection, despite the establishment of a productive infection (Fig. S1B-I). To gain insight into the kinetics of vtRNA expression during viral infection, we performed a time course experiment in A549 cells infected with EMCV (Fig. 1H) and HEK293 cells infected with SINV (Fig. 1I). In both cases, upregulation of vtRNA expression occurred rather late during the course of viral infection. VtRNA expression was noticeably increased at 24h (EMCV) and 32h (SINV) post-infection and continued to rise at least until 48h (EMCV) or 72h (SINV). Upregulation of vtRNAs was also dose-dependent (Fig. S1J).

**Figure 1:**
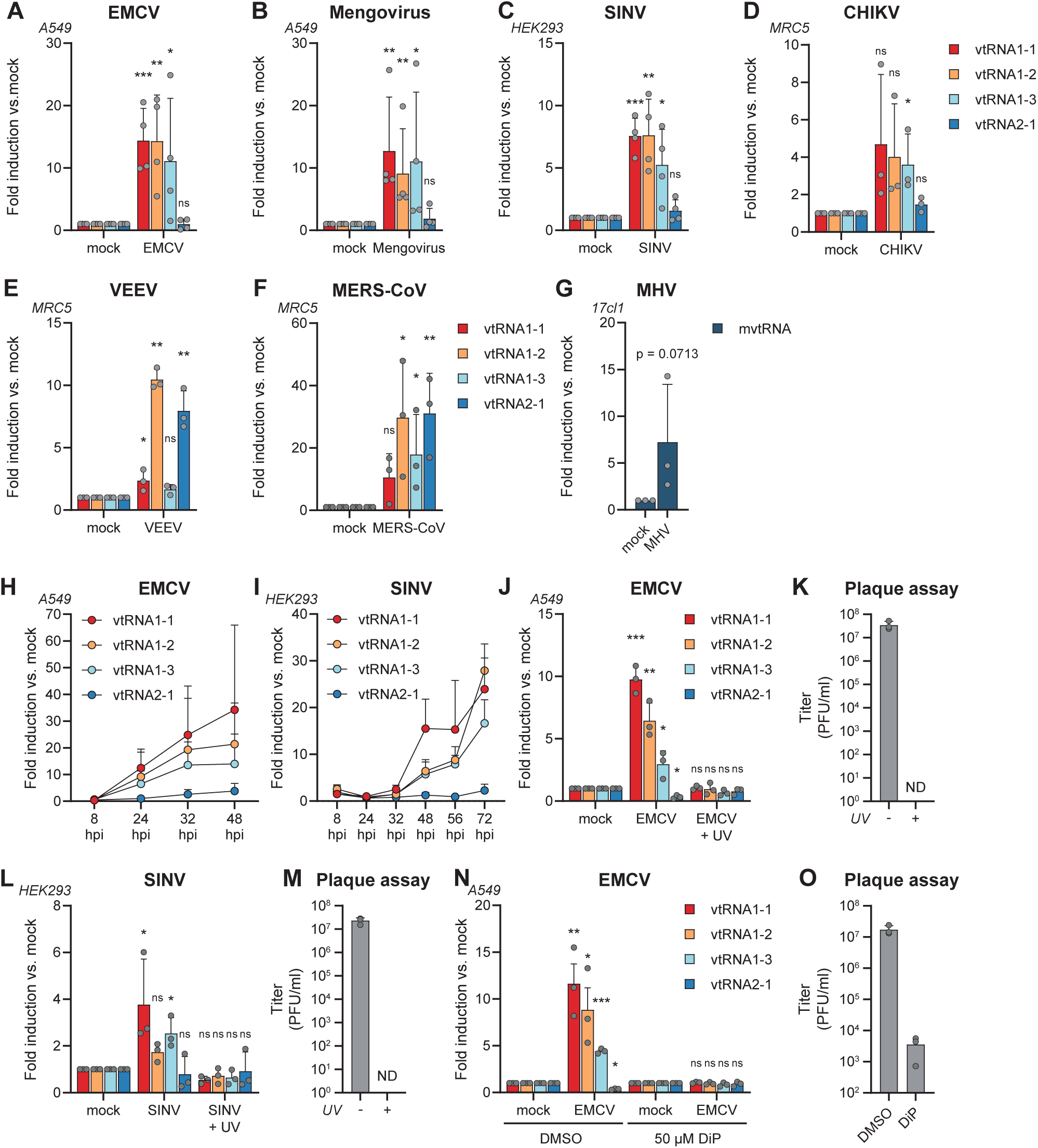
A diverse repertoire of RNA viruses induces increased vtRNA expression in a replication-dependent manner. **A)** A549 cells were infected with EMCV for 24h at MOI = 0.25. Expression of vtRNA1-1, vtRNA1-2, vtRNA1-3, and vtRNA2-1 was measured by RT-qPCR and normalized against expression of a housekeeping gene (*RN18S1*). Values are displayed relative to mock-infected cells (n=4). **B)** A549 cells were infected with Mengovirus for 24h at MOI = 0.5. Expression of vtRNAs was measured by RT-qPCR as in (A). Values are displayed relative to mock-infected cells (n=4). **C)** HEK293 cells were infected with GFP-tagged Sindbis virus for 48h at MOI = 1. Expression of vtRNAs was measured by RT-qPCR as in (A). Values are displayed relative to mock-infected cells (n=4). **D)** MRC5 cells were infected with Chikungunya virus (CHIKV) for 24h at MOI = 1. The upregulation of vtRNA expression was measured by RT-qPCR as in (A). Values are displayed relative to mock-infected cells (n=3). **E)** MRC5 cells were infected with Venezuelan Equine Encephalitis Virus (VEEV) for 48h at MOI = 1. The upregulation of vtRNA expression was measured by RT-qPCR as in (A). Values are displayed relative to mock-infected cells (n=3). **F)** MRC5 cells were infected with Middle East respiratory syndrome coronavirus (MERS-CoV) for 48h at MOI = 1. The upregulation of vtRNA expression was measured by RT-qPCR as in (A). Values are displayed relative to mock-infected cells (n=3). **G)** The mouse cell line 17cl1 was infected with mouse hepatitis virus (MHV) for 16h at MOI = 1. The expression level of mouse vtRNA (mvtRNA) was determined by RT-qPCR. Expression levels were normalized against mouse *Rn18s1* and shown relative to mock-infected cells (n=3). **H)** Kinetics of EMCV-induced vtRNA upregulation in A549 cells. Cells were infected with EMCV and harvested at the indicated time points post-infection. Expression levels of vtRNAs were monitored by RT-qPCR and normalized as in (A). Data are displayed as fold induction over the mock-infected condition of each time point (n=3). **I)** Kinetics of SINV-induced vtRNA upregulation in HEK293 cells. Cells were infected with SINV and were harvested at the indicated time points post-infection. Expression levels of vtRNAs were monitored by RT-qPCR and normalized as in (A). Data are displayed as fold induction over the mock-infected condition of each time point (n=3). **J)** A549 cells were infected with live or UV-irradiated EMCV. 24hpi, cells were harvested and vtRNA expression levels were assessed by RT-qPCR. Data were normalized as in (A). Data are displayed as fold induction over the mock-infected condition (n=3). **K)** EMCV titers in (J) were determined by plaque assay (n=3). **L)** HEK293 cells were infected with live or UV-irradiated SINV. Cells were harvested 48hpi and vtRNA expression levels were assessed by RT-qPCR. Data were normalized as in (A). Data are displayed as fold induction over the mock-infected condition (n=3). **M)** SINV titers in (L) were determined using plaque assay (n=3). **N)** A549 cells were treated with dipyridamole (DiP), an inhibitor of cardiovirus replication, or a vehicle control (DMSO). After 30 min of pretreatment, cells were infected with EMCV, or left uninfected. Cells were harvested 24hpi and vtRNA expression levels were assessed by RT-qPCR. Data were normalized as in (A). Data are displayed as fold induction over the mock-infected cells (n=3). **O)** EMCV titers in (N) were determined using plaque assay (n=3). Data are means ± s.d.. Statistical analyses in Fig. 1A-G, 1J, 1L and 1N were performed on unnormalized, log-transformed data using a Student’s *t*-test to test significance of each vtRNA expression between virus-infected and mock-infected conditions.

We conclude that infection with a broad range of RNA viruses enhances vtRNA expression in a time- and dose-dependent manner in human and murine cells, but that vtRNA upregulation is not a general feature of the host response to all viral pathogens.

### VtRNA upregulation is independent of type I IFN signaling and requires active viral replication

Next, we explored how such a broad range of viruses can upregulate vtRNAs. We reasoned that there must be a common denominator that drives vtRNA expression during these diverse infections. We found that vtRNA expression is not increased upon treatment with type I IFNs (Fig. S1K-L), in line with a previous report^39^. In addition, transfection of HEK293 cells with poly(I:C), a double-stranded RNA mimic that activates RIG-I-like receptors (RLRs) and the type I IFN response, did not increase vtRNA expression (Fig. S1M-N). Since vtRNA expression increases over time (Fig. 1H-I), we hypothesized that active viral replication is required to drive vtRNA upregulation. Indeed, in contrast to cells infected with live EMCV, the expression of vtRNAs remained unchanged after exposure of cells to UV-inactivated EMCV (Fig. 1J). We confirmed successful UV-inactivation by plaque assay 24h post-infection (Fig. 1K). Similarly, UV-inactivation of SINV abolished vtRNA upregulation (Fig. 1L-M). This suggests that viral replication is required for vtRNA upregulation. To further test this, we infected cells with EMCV in the presence or absence of the cardiovirus replication inhibitor dipyridamole (DiP). Treatment with DiP strongly inhibited EMCV replication, which prevented vtRNA upregulation (Fig. 1N-O). Altogether, these results indicate that ongoing viral replication is required to drive vtRNA upregulation.

### Loss of vtRNAs impairs replication of SINV

Given the broad range of viral infections that induce vtRNA expression (Fig. 1), we speculated that vtRNAs either support the antiviral host response or are co-opted by viruses to aid their replication. To investigate this, we used CRISPR-Cas9 genome engineering to generate HEK293 cells that lack all three vtRNA1 genes (vtRNA1-1, vtRNA1-2, and vtRNA1-3), hereafter referred to as vtRNA1 KO. Two universal guide RNAs were designed that simultaneously target the two highly homologous internal promoter boxes in the *VTRNA1-1*, *VTRNA1-2,* and *VTRNA1-3* genes on chromosome 5 (Fig. 2A). Because vtRNA2-1 expression remained unchanged during most viral infections, we left the *VTRNA2-1* locus intact. Five vtRNA1 KO clones were generated. PCR analysis using primers flanking each vtRNA gene demonstrated precise excision of the vtRNA genes from the genome (clones 1, 2, and 3) or a more extensive disruption of the vtRNA1 locus (clones 4 and 5) (Fig 2B). RT-qPCR analysis demonstrated that all five clones lack detectable vtRNA1-1, vtRNA1-2, and vtRNA1-3 expression, while vtRNA2-1 expression is unaffected (Fig. 2C). Northern blotting further confirmed the absence of vtRNA1-1, vtRNA1-2, and vtRNA1-3 expression (Fig. 2D). We proceeded to infect wild-type (WT) cells and vtRNA1 KO clones with the prototype alphavirus SINV to determine how loss of vtRNAs impacts SINV replication. Importantly, all five vtRNA1 KO clones consistently showed a significant decrease in viral titer and viral transcripts at 48h post-infection compared to WT cells (Fig. 2E-F). To validate that impaired SINV replication was due to loss of vtRNAs, we reintroduced vtRNA1 expression in three vtRNA1 KO clones by means of lentiviral transduction using individual vectors encoding vtRNA1-1, vtRNA1-2, or vtRNA1-3, or, as a control, empty vectors. Following antibiotic selection, vtRNA1-1 and vtRNA1-2 expression was restored to WT levels in these cell lines. vtRNA1-3 expression was also restored, yet remained 4- to 7-fold lower compared to WT cells (Fig. 2G-I). When infecting these cells with SINV, viral replication was doubled in all three rescued cell lines compared to the vtRNA1 KO cell lines transduced with an empty vector (Fig. 2J-L). Note that, unlike endogenous vtRNAs, the expression of vtRNA transgenes is not upregulated in response to viral infection (Fig. 2G-I , which may explain why viral replication is increased but not fully restored. Altogether, these data indicate that SINV replication is impaired in vtRNA1 KO cells and that this is rescued by reintroduction of vtRNAs.

**Figure 2:**
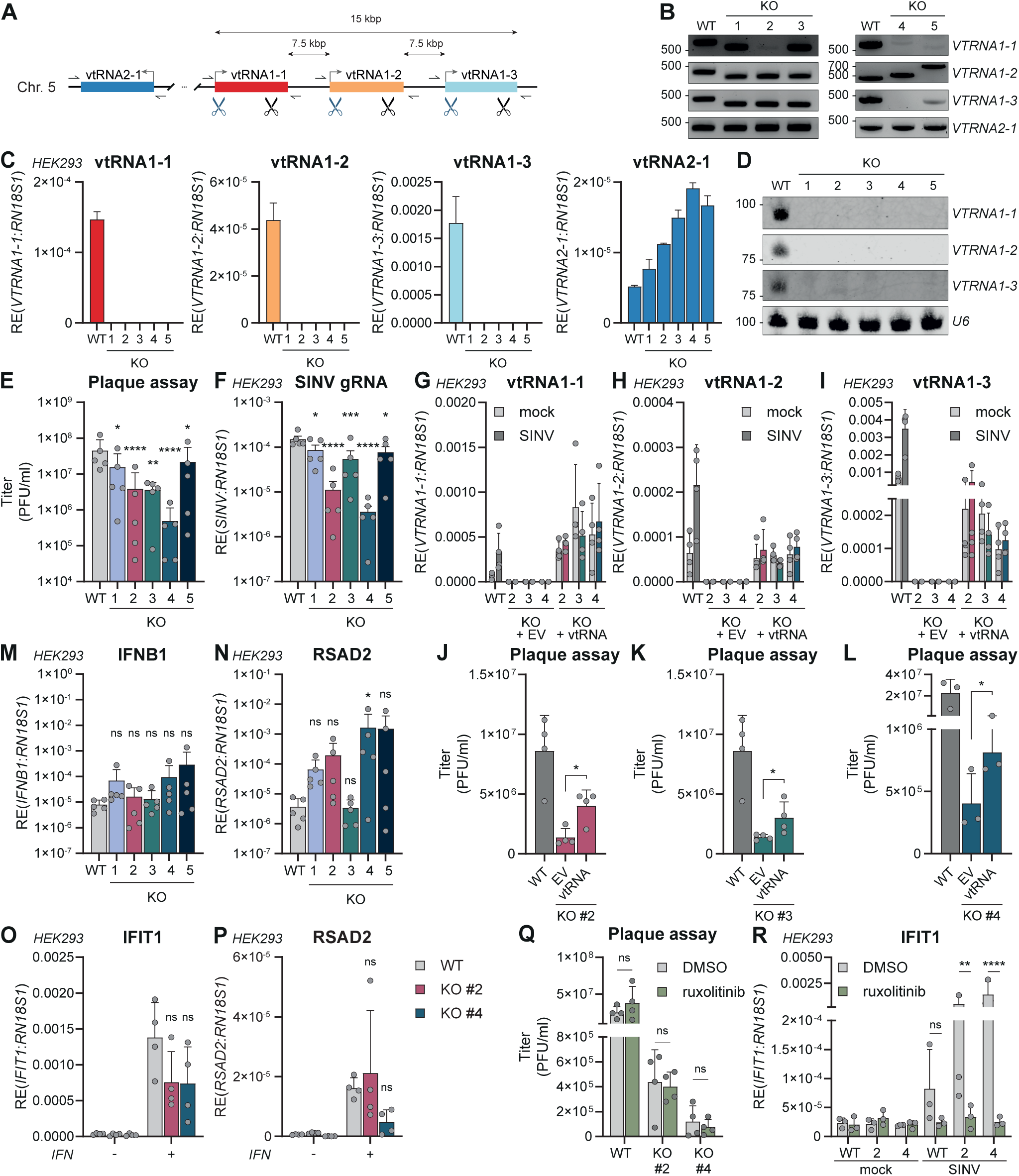
SINV replication is inhibited in vtRNA1 KO cells in a type I interferon-independent manner. **A)** Schematic overview of the CRISPR-Cas9 genome editing strategy to knockout *VTRNA1-1*, *VTRNA1-2* and *VTRNA1-3*. Two single guide RNAs were designed that simultaneously target regions in the *VTRNA1-1*, *VTRNA1-2* and *VTRNA1-3* genes. Primer sites used in (B) are indicated with half arrows. **B)** Primers flanking each *VTRNA* gene were designed to assess the integrity of each *vtRNA* gene after CRISPR-Cas9 genome editing, compared to HEK293 WT cells. Five monoclonal cell lines (1 to 5) were chosen based on disruption of the *VTRNA* loci. One representative experiment is shown (n=2). **C)** Loss of vtRNA expression in the clones selected in (B) was shown by RT-qPCR. Expression of vtRNAs was measured and normalized to *RN18S1*. One representative experiment is shown (n=4). **D)** Northern Blot showing the loss of vtRNA1-1, vtRNA1-2, and vtRNA1-3 expression in the clones selected in (B). U6 snRNA expression is shown as housekeeping control. **E)** HEK293 WT or vtRNA1 KO cell lines were infected with SINV at MOI = 1. Viral replication was assessed by plaque assay 48hpi (n=5). **F)** RT-qPCR showing the relative expression of SINV genomic RNA (‘gRNA’) in cells infected in (E), using primers directed against the *nsP1* gene in the SINV genome. Data are normalized against *RN18S1*. (n=5). **G)** VtRNA1 KO cells were transduced with empty lentiviruses (+ EV) or lentiviruses encoding *VTRNA1-1*, *VTRNA1-2*, and *VTRNA1-3* (+ vtRNA). Expression of vtRNA1-1 in mock-infected or SINV-infected WT HEK293 and rescued vtRNA1 KO cells was analyzed by RT-qPCR 48hpi (n=4). **H)** Expression of vtRNA1-2, as in (G) (n=4). **I)** Expression of vtRNA1-3, as in (G) (n=4). **J)** Replication of SINV 48hpi in WT, EV-transduced, or vtRNA1-transduced KO clone 2 was measured using plaque assay (n=4). **K)** Replication of SINV 48hpi in WT, EV-transduced, or vtRNA1-transduced KO clone 3 was measured using plaque assay. WT replication levels are the same as in (J) (n=4). **L)** Replication of SINV 72hpi in WT, EV-transduced, or vtRNA1-transduced KO clone 4 was measured using plaque assay (n=3). **M)** Relative expression of IFN-β mRNA was measured by RT-qPCR, normalized as in (F) (n=5). **N)** Relative expression of RSAD2 mRNA was measured by RT-qPCR, normalized as in (F) (n=5). **O-P)** HEK293 WT, vtRNA1 KO clone 2 or clone 4 were stimulated with exogenous type I IFN for 24h. Expression of IFIT1 and RSAD2 mRNA was measured by RT-qPCR, and was normalized as in (F) (n=4). **Q)** SINV replication in HEK293 WT, vtRNA1 KO clone 2 or clone 4 48hpi, as measured by plaque assay. Cells were treated with ruxolitinib or vehicle control (DMSO) before and during the infection (n=4). **R)** Expression of IFIT1 mRNA in mock-infected or SINV-cells in (Q), measured using RT-qPCR. Data was normalized as in (F) (n=3). Data are represented as means ± s.d.. Statistical analyses in 2E-F, M-N and 2O-R were performed using a repeated measures one-way ANOVA on unnormalized, log-transformed data. Data were compared with the WT control. In 2J-L, a Student’s *t*-test was performed on log-normalized data, between EV- and vtRNA1-transduced cells.

### VtRNAs do not modulate the type I IFN response during SINV infection

Next, we investigated whether vtRNAs impact viral replication via modulation of the antiviral type I IFN response. We first tested whether vtRNA1 KO cells differentially respond to treatment with recombinant type I IFN using RT-qPCR analysis of transcripts encoding IFIT1 and RSAD2 (two prototype IFN-stimulated genes (ISGs)). Upregulation of IFIT1 and RSAD2 expression was comparable across WT cells and all vtRNA1 KO clones, indicating that vtRNA1-deficient cells do not have an intrinsic defect in type I IFN signaling (Fig. 2O-P). We then analyzed the antiviral type I IFN response in SINV-infected WT and vtRNA1 KO cells by RT-qPCR analysis of IFN-β and RSAD2 transcripts. SINV-induced IFN-β expression was not significantly affected by loss of vtRNAs, whereas RSAD2 expression was elevated in some, but not all, vtRNA1 KO clones (Fig. 2M-N). To understand whether this increase in ISG expression, although inconsistent across clones, is responsible for the decrease in SINV replication (Fig. 2E-F), we pretreated WT and vtRNA1 KO cells with the JAK1/JAK2-inhibitor ruxolitinib prior to SINV infection. Ruxolitinib inhibits signaling events downstream of the type I interferon receptor (IFNAR) and thus negates the antiviral effect of IFN-β expression. Replication of SINV in the vehicle-treated vtRNA1 KO cells was strongly decreased compared to WT HEK293 cells (Fig. 2Q), as observed before (Fig. 2E-F). Treatment of WT cells with ruxolitinib had a negligible effect on SINV replication (Fig. 2Q) despite potently suppressing ISG induction (Fig. 2R) indicating that under the conditions tested, the type I IFN response does not play a major role in restricting SINV. Importantly, SINV replication was unaltered in ruxolitinib-treated vtRNA1 KO cells compared to untreated vtRNA1 KO cells (Fig. 2Q), indicating that the increased ISG response in some vtRNA1 KO clones is not accountable for the reduction in SINV replication (Fig. 2E). We speculate that the increase in ISG expression observed in these vtRNA1 KO clones may therefore be a consequence (rather than a cause) of decreased SINV replication and the reduced accumulation of virus-encoded IFN antagonists that inhibit JAK-STAT signaling downstream of the IFNAR^43–45^. Taken together, we conclude that the proviral function of vtRNA1-1, vtRNA1-2, and vtRNA1-3 in SINV infection does not involve modulation of the type I IFN response.

### Loss of vtRNAs impairs replication of EMCV and Mengovirus

To address whether the loss of vtRNAs similarly impairs infection with a virus unrelated to SINV, we used the CRISPR-Cas9 procedure described above in A549 cells to generate a monoclonal vtRNA1 KO cell line. Correct targeting of the vtRNA1 locus was confirmed by genomic PCR, RT-qPCR, and Northern blot (Fig. S2A-C). As observed for SINV, infection of vtRNA1 KO A549 cells with EMCV or Mengovirus resulted in decreased viral titers (Fig. S2D and Fig. S2P) and decreased viral transcription (Fig. S2E) compared to A549 WT, which could not be explained by an altered type I IFN response (Fig. S2F-G). We corroborated these results using an antisense oligonucleotide (ASO)-mediated approach to simultaneously knockdown vtRNA1-1, vtRNA1-2, and vtRNA1-3. Depletion of vtRNA1-1, vtRNA1-2, and vtRNA1-3 impaired replication of EMCV, compared to cells treated with non-targeting control ASOs (Fig. S2H-L). We also restored vtRNA1-1, vtRNA1-2, and vtRNA1-3 expression in the vtRNA1 KO A549 cells by means of lentiviral transduction, and infected these cells with Mengovirus (Fig. S2M-O).

The decrease in viral titer observed in vtRNA1 KO A549 cells was restored upon reintroduction of vtRNAs (Fig. S2P). Altogether, we conclude that vtRNAs are co-opted by SINV, EMCV, and Mengovirus to sustain their replication and that vtRNAs have a proviral function.

### Identification of the vtRNA1 interactome by RNA affinity purification and proteomics

To understand how vtRNAs affect viral replication, we used RNA antisense purification coupled to mass spectrometry (RAP-MS) to identify proteins that interact with vtRNAs^46^. In RAP-MS, intact cells are irradiated with UV-C light (254 nm) prior to cell lysis to create covalent bonds between RNA molecules and interacting proteins. The vtRNA-protein complexes are retrieved from cellular lysates under highly denaturing conditions using biotinylated hybridizing DNA oligonucleotides specific for vtRNAs. This allows the isolation of native vtRNAs and the identification of proteins interacting with vtRNAs by mass spectrometry (Fig. 3A). To capture vtRNAs, we designed three individual 80 nt-long DNA probes that are reverse complementary to vtRNA1-1, vtRNA1-2, or vtRNA1-3 (named ‘antisense’). As negative control, we used the same targeted region in the vtRNA but used the coding (‘sense’) sequence. The probes specific for vtRNA1-1, vtRNA1-2, and vtRNA1-3 pulled down 40, 69, and 65 unique proteins over negative controls, respectively. We noted that the interactomes of individual vtRNAs have substantial overlap, but also contain unique interacting proteins (Fig. 3B). In all pulldowns, we retrieved telomerase-associated protein 1 (TEP1), which is responsible for docking vtRNAs to the vault complex^13–15^. We also identified Sjögren syndrome type B antigen (SSB), another known interactor of vtRNAs^47^, further confirming the validity of our screen. Next, we performed a Gene Ontology enrichment analysis of the interactors that were identified in at least two of the three vtRNA interactomes to gain insight into the biological processes in which vtRNA interactors are involved. The GO Term “Regulation of mRNA metabolic process” was the most enriched biological process amongst the shared interactors of the three vtRNAs (Fig. 3C). Similar terms were found in GO Term analyses of enriched interactors of each separate vtRNA (Fig. S3A-C). Consistently, many of the enriched interactors are known or predicted RBPs (Fig. 3D-F, highlighted in blue). On the other hand, no proteins related to type I IFN induction or IFNAR signaling were identified, not even when the RAP-MS screen was repeated in EMCV-infected cells (Fig. S3D-F), further suggesting that vtRNAs do not directly modulate the antiviral type I IFN response, at least in the context of EMCV infection.

**Figure 3:**
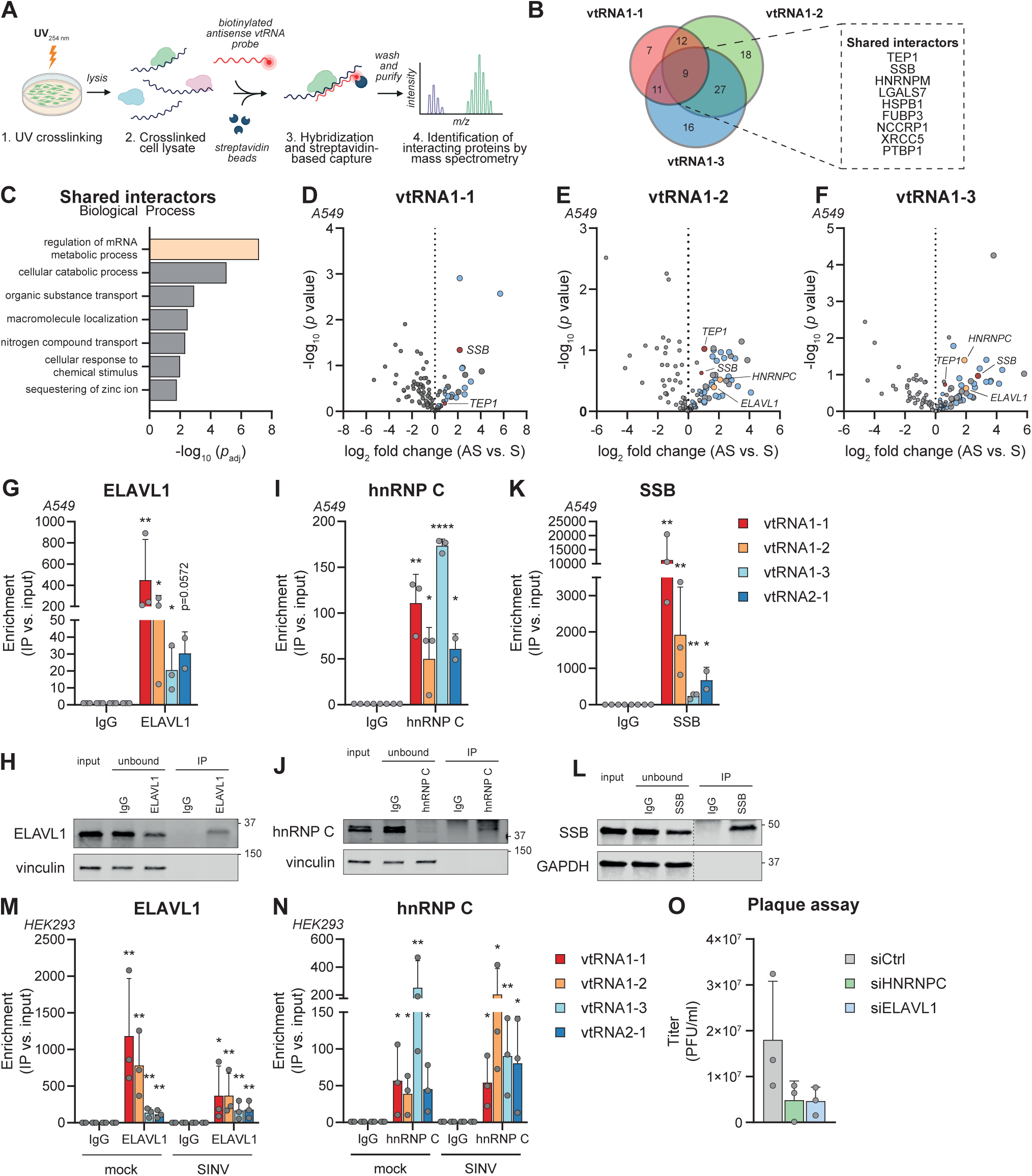
A RAP-MS screen uncovers the vtRNA1 interactome, including interactions with RNA-binding proteins hnRNP C and ELAVL1. **A)** Schematic representation of the RAP-MS pipeline. A549 cells are irradiated with UV-C light (254 nm) to generate covalent bonds between interacting RNAs and proteins. After lysis, vtRNA-protein complexes are retrieved using hybridizing biotinylated DNA probes and streptavidin-coated magnetic beads. After stringent washing steps, isolated proteins are identified using LC-MS/MS. **B)** Venn diagram showing the overlap in the interactomes of vtRNA1-1, vtRNA1-2 and vtRNA1-3. The panel on the right highlights the 9 proteins that were retrieved in all three interactomes. **C)** GO Term analysis showing the enriched Biological Processes of interactors that were retrieved in at least two of the three vtRNA interactomes. **D-F)** Quantification of vtRNA-interacting proteins (antisense, ‘AS’) compared to the negative control (sense, ‘S’), expressed as log_2_-fold change. The enriched proteins that are classified as “RNA-binding” according to GO Term analysis are highlighted in blue. Positive controls are highlighted in red, and proteins of interest are highlighted in yellow. Statistical analysis was performed using a two-tailed *t*-test with a *post-hoc* FDR correction (n=2). **G)** RIP-qPCR showing the interaction between vtRNAs and ELAVL1. Immunoprecipitations were performed on A549 cell lysates with anti-ELAVL1 or mouse IgG1 antibodies. Coprecipitating vtRNAs in the immunoprecipitates were quantified using RT-qPCR and expressed as fraction of input vtRNA levels. Enrichment of vtRNAs was then calculated by normalization to the IgG control (n=3). **H)** The efficiency of ELAVL1 immunoprecipitation, described in (G), was determined by immunoblotting. Vinculin staining serves as a housekeeping marker to show equal loading. A representative image is shown (n=3). **I)** RIP-qPCR showing the interaction between vtRNAs and hnRNP C in A549 cells, performed as in (G) (n=3). **J)** The efficiency of hnRNP C immunoprecipitation, described in (I), was determined by immunoblotting. Vinculin staining serves as a housekeeping marker to show equal loading. Representative image is shown (n=3). **K)** RIP-qPCR showing the interaction between vtRNAs and SSB in A549 cells, performed as in (G) (n=3). **L)** The efficiency of SSB immunoprecipitation, described in (K), was determined by immunoblotting. Vinculin staining serves as a housekeeping marker to show equal loading. Dotted line represents the juxtaposition of two non-adjacent lanes on the same membrane. Representative image is shown (n=3). **M)** RIP-qPCR showing the enrichment of vtRNAs on ELAVL1 in mock- and SINV-infected HEK293 cells. The co-precipitating vtRNAs were expressed relative to the input expression levels and normalized to the enrichment in the IgG isotype control IP performed on either mock- or SINV-infected lysates (n=3). **N)** RIP-qPCR showing the enrichment of vtRNAs on hnRNP C in mock- and SINV-infected HEK293 cells. Enrichments were calculated as in (M) (n=3). **O)** SINV titers in HEK293 cells depleted of hnRNP C or ELAVL1 by means of siRNAs. Cells were treated with siRNAs for 48h and then infected with SINV. Titers were determined using plaque assay 48hpi (n=3). Data in bar graphs are represented as means ± s.d.. Statistical analysis in 3G, 3I, 3K, and 3M-N was performed using a one-sample *t*-test for each vtRNA compared to the IgG control on log-transformed enrichment values.

### VtRNAs interact with the RNA binding proteins hnRNP C and ELAVL1

Among the identified vtRNA1-interacting proteins with a role in regulating RNA metabolic processes are the proteins ELAVL1 (also known as HuR) and hnRNP C (Fig 3E-F). Interactions with these RNA binding proteins persist upon EMCV-infection (Fig S3D-F), indicating that these interactions may play a role during infection. Additionally, ELAVL1 interacts with vtRNA2-1 in intestinal epithelial cells^30^. We carried out RNA immunoprecipitations coupled to RT-qPCR (RIP-qPCR) to validate our findings.

ELAVL1 and hnRNP C were immunoprecipitated from A549 cells and the presence of vtRNAs amongst the co-precipitating RNA was analyzed by RT-qPCR. All four vtRNAs were strongly enriched in the hnRNP C and ELAVL1 eluates compared to an isotype-matched control IgG IP (Fig. 3G-J). As a positive control, we showed strong interactions between SSB and all four vtRNAs (Fig. 3K-L). ELAVL1 interacts most strongly with vtRNA1-1, while hnRNP C interacts more strongly with vtRNA1-3. This may point to a preferential affinity for a specific vtRNA. Next, we investigated whether interactions between either ELAVL1 or hnRNP C and vtRNAs are altered during SINV infection. In mock-infected HEK293 cells, we found an almost identical interaction pattern between vtRNAs and ELAVL1 or hnRNP C as observed in A549 cells (Fig. 3M-N, Fig. S3G-H). During SINV infection, these interactions persisted.

SiRNA-mediated depletion of ELAVL1 or hnRNP C inhibited SINV replication in HEK293 cells (Fig. 3O, Fig. S3I), as previously reported for ELAVL1 by others^48^. Loss of ELAVL1 or hnRNP C thus phenocopies the loss of vtRNA1, potentially placing each of these factors in the same proviral pathway. In conclusion, we uncovered interactions between vtRNAs and ELAVL1 and hnRNP C at homeostatic conditions that persist during viral infection.

### vtRNAs are required for nuclear export of ELAVL1 and hnRNP C upon infection

In uninfected cells, ELAVL1 shuttles between the nucleus and cytoplasm, whereas hnRNP C is mostly located in the nucleus^49,50^. Both proteins translocate to the cytoplasm during infection with SINV or other alphaviruses, where they interact with SINV RNA and aid viral replication^48,51–55^. For example, ELAVL1 stabilizes SINV RNA and protects it against RNA decay^48^. In addition, both proteins promote replication of diverse viruses, including hepatitis B virus, MERS-CoV, and poliovirus^56–59^. Thus, hnRNP C and ELAVL1 are hijacked by viruses and are important for optimal viral replication, similar to vtRNAs (Fig. 2 and Fig. S2). We therefore set out to discover whether and how vtRNAs are required for the proviral activity of ELAVL1 and hnRNP C during SINV and EMCV infection.

To test whether vtRNAs affect nuclear-cytoplasmic shuttling of ELAVL1 and hnRNP C, we set up a cell fractionation assay. We infected WT HEK293 and two vtRNA1 KO clones with SINV for 48h and subsequently fractionated the cells into three fractions containing cytoplasmic, membranous, or nuclear proteins. In mock-infected WT and vtRNA1 KO cells, hnRNP C and ELAVL1 predominantly localized to the nuclear compartment (Fig. 4A-C). Upon SINV infection, hnRNP C and ELAVL1 relocated to the membrane-containing and cytoplasmic fractions in WT cells (Fig. 4A-C). Instead, the nuclear export of hnRNP C and ELAVL1 was strongly diminished in SINV-infected vtRNA1 KO cells (Fig. 4A-C). We visualized the altered nuclear-cytoplasmic transport of ELAVL1 in SINV-infected vtRNA1 KO cells at single-cell level using a microscopy-based approach. We used an antibody directed against dsRNA to identify SINV-infected cells and assessed the localization of ELAVL1 in dsRNA-positive cells. Consistent with the fractionation experiments, ELAVL1 relocated to the cytoplasm in SINV-infected WT cells, whereas it remained in the nucleus in SINV-infected vtRNA1 KO cells (Fig. 4D-E). To test whether vtRNAs generally impact nucleocytoplasmic transport, we assessed the subcellular localization of other proteins. The localization of the predominantly cytoplasmic vtRNA-binding protein SSB or the unrelated nuclear protein MORC3 was unaffected by the loss of vtRNAs, both in uninfected and SINV-infected cells (Fig. S4A-C). In addition, we assessed the relocalization of the RNA helicase AQR, which was not identified in the RAP-MS screen as a vtRNA interactor yet is shuttled out of the nucleus upon SINV infection^52^ In absence of vtRNA1 expression, AQR was still exported from the nucleus following viral infection in vtRNA1 KO clone 2 (Fig. S4A and D). Together, these findings suggest that vtRNAs specifically aid the nuclear export of ELAVL1 and hnRNP C during SINV infection, without generally affecting nucleocytoplasmic protein distribution. We corroborated these findings in A549 WT and vtRNA1 KO cells infected with EMCV. Consistent with the observations described above, ELAVL1 and hnRNP C are predominantly located in the nucleus in uninfected WT and vtRNA1 KO A549 cells (Fig. S4E-G, I). Both proteins translocate to the cytoplasm upon EMCV infection in WT cells, which was less evident in vtRNA1 KO cells (Fig. S4E-G). Instead, the localization of SSB was unaltered by vtRNA expression in both uninfected and EMCV-infected cells (Fig. S4E and H). Altogether, we conclude that vtRNAs facilitate the translocation of ELAVL1 and hnRNP C from the nucleus to the cytosol during both SINV and EMCV infection.

**Figure 4:**
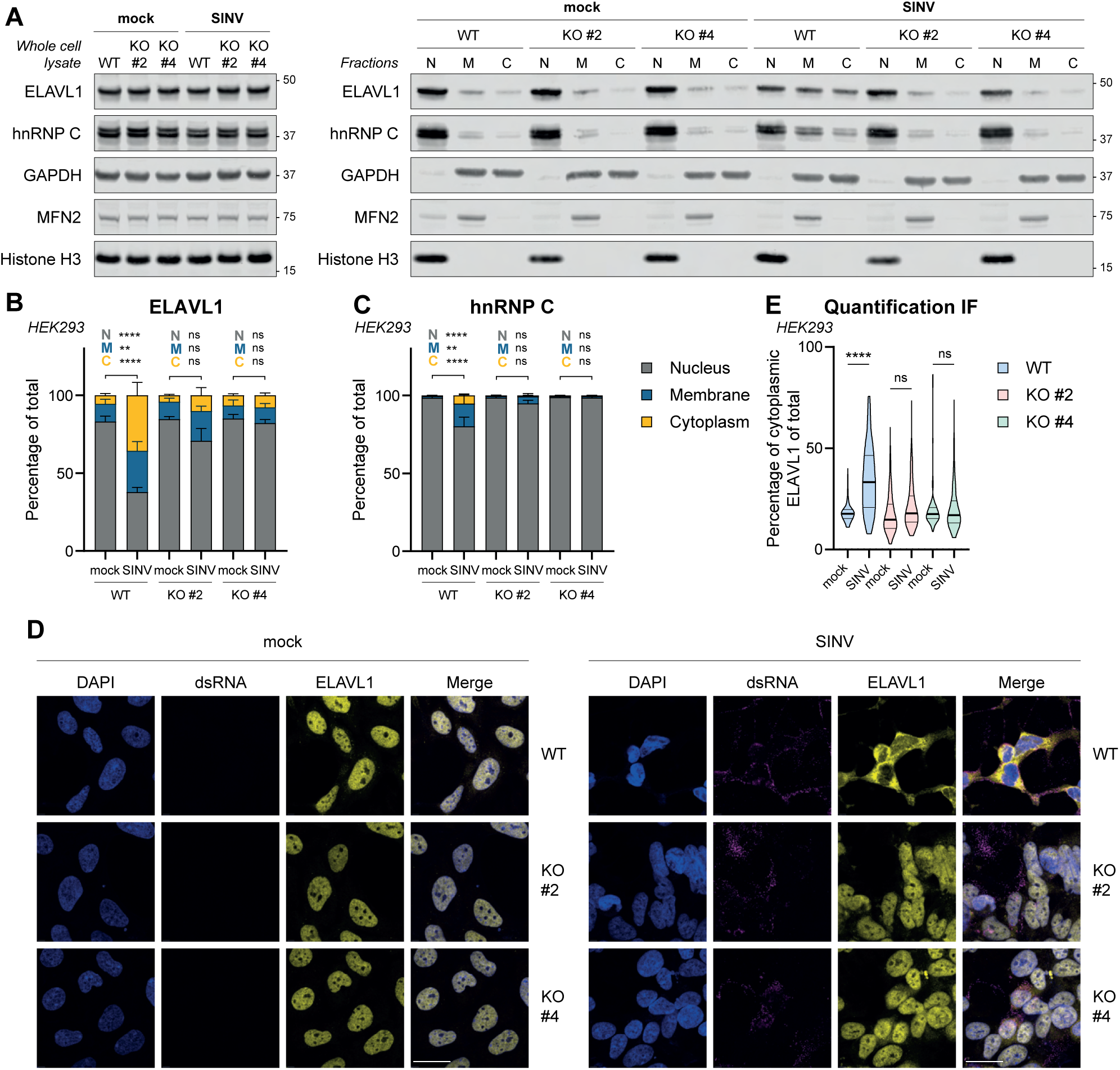
vtRNA1s promote ELAVL1 and hnRNP C nuclear export during SINV infection. **A)** Expression of ELAVL1 and hnRNP C in whole cell lysate (left) or in different subcellular compartments (right). Cell lysates of uninfected or SINV-infected HEK293 WT or vtRNA1 KO cells (clone 2 and 4) were fractionated into a nuclear (N), membranous (M), and cytoplasmic (C) fraction. GAPDH, mitofusin 2 (MFN2), and histone H3 are used as markers for the cytoplasmic/membranous, membranous, and nuclear fraction, respectively. One representative experiment is shown (n=3). **B-C)** Quantification of ELAVL1 and hnRNP C band intensities in (A). Intensity of each band was expressed as a percentage of cumulative intensity of the three fractions within each condition. Data represent means ± s.d. (n=3); *p* values for statistical tests between every fraction are shown preceding with ‘N’ (nucleus), ‘M’ (membrane), or ‘C’ (cytoplasm). **D)** HEK293 WT or vtRNA1 KO cells (clone 2 or 4) were plated in chamber slides for immunofluorescence microscopy. Cells were fixed 48hpi with SINV and stained for ELAVL1 (yellow) and dsRNA (magenta). Nuclei were stained with DAPI (blue). Scale bar indicates 25 µm. Representative micrographs are shown (n=3). **E)** Quantification of ELAVL1 staining intensity in the cytoplasm of cells described in (D). Segmentation of cells into nuclei and cytoplasm was done based on DAPI and phalloidin staining. Only cells showing dsRNA staining as a marker of infection were included in the analysis. The truncated violin plot indicates the median (central, thick line), interquartile range (two thinner lines), and the minimum and maximum values of each distribution. At least 120 cells per condition were quantified. A representative experiment is shown (n=2). Statistical analysis in 4B-C was performed using a repeated measures one-way ANOVA on unnormalized, log-transformed data. In 4E, analysis was performed using a regular one-way ANOVA.

### Loss of vtRNAs interferes with the proviral activity of hnRNP C and ELAVL1

To test whether hnRNP C and ELAVL1 require the presence of vtRNAs to execute their proviral activities, we treated WT or vtRNA1 KO cells with siRNAs targeting hnRNP C or ELAVL1 for 48h, followed by SINV infection. Depletion of hnRNP C and ELAVL1 both significantly diminished viral replication in WT cells (Fig. 5A-B), consistent with our previous observations (Fig. 3O). Instead, in vtRNA1-deficient cells, siRNA-mediated depletion of hnRNP C and ELAVL1 led to a smaller reduction in viral replication (Fig. 5A-B). This was most evident for hnRNP C; depletion of hnRNP C reduced SINV replication to 46% in WT cells versus 82% in vtRNA1 KO cells. This suggests that hnRNP C requires the presence of vtRNAs to mediate its proviral effects. In the case of ELAVL1, this effect was less pronounced, as loss of ELAVL1 reduced SINV replication to 16% in WT cells versus 31% in vtRNA1 KO cells (Fig. 5A-B). Possibly, the proviral activity of ELAVL1 is less affected by the loss of vtRNAs because, unlike hnRNP C, ELAVL1 already localizes to some degree in the cytoplasm in uninfected cells regardless of vtRNA expression (Fig. 4A-C). These findings suggest that the proviral activity of hnRNP C and, to a lesser extent of ELAVL1, requires the presence of vtRNAs.

**Figure 5:**
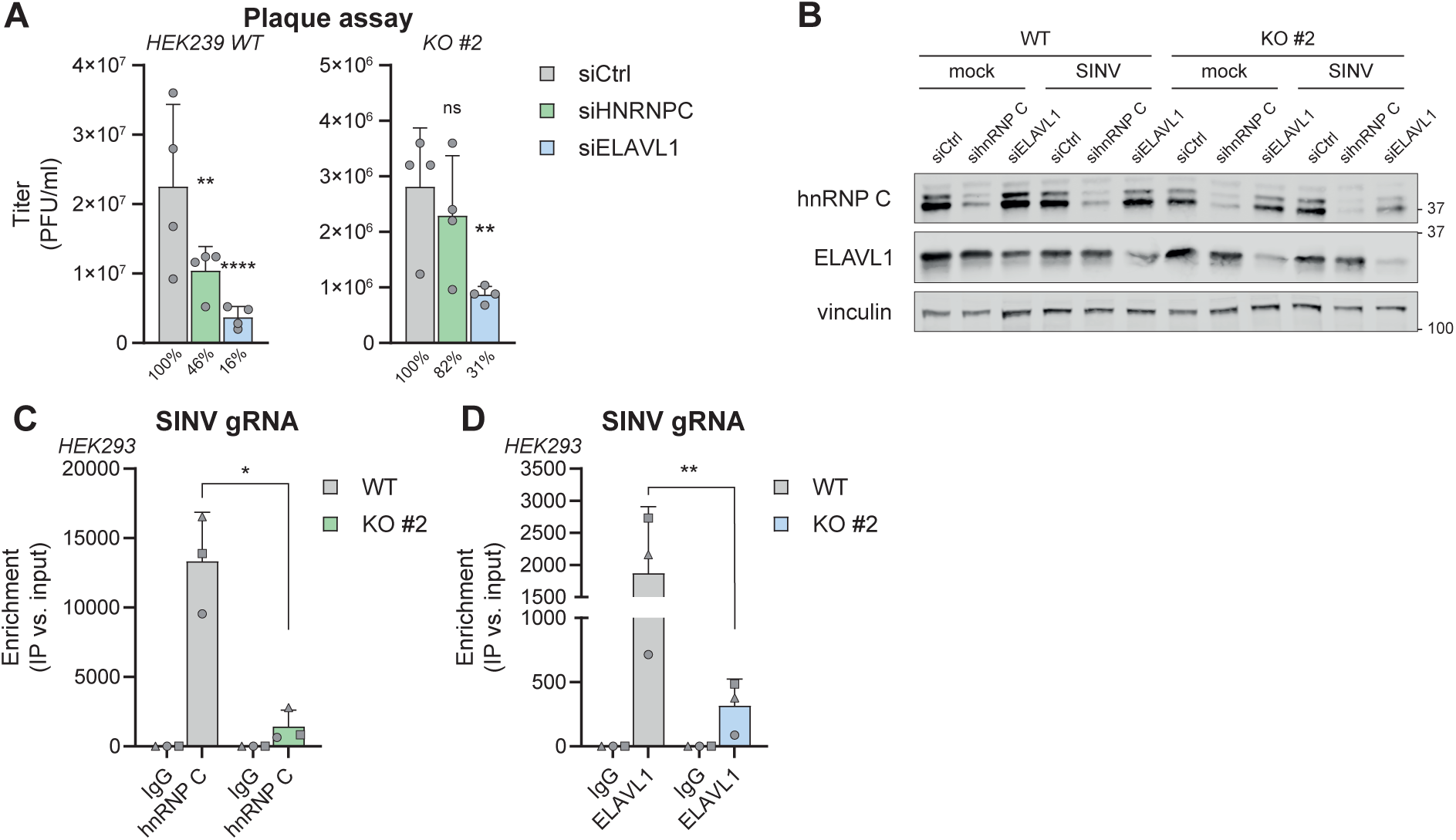
ELAVL1 and hnRNP C depletion inhibits SINV replication in a vtRNA1-dependent manner. **A)** SINV titers in HEK293 WT cells or vtRNA1 KO clone 2 following siRNA-mediated depletion of hnRNP C or ELAVL1. Cells were treated with siRNAs for 48h and subsequently infected with SINV for 48h. Percentages below bars represent the mean relative decrease in SINV titer compared to the siCtrl-condition (n=4). **B)** Validation of the knockdown of hnRNP C or ELAVL1 in (A) and (B) by immunoblotting. Vinculin staining serves as a housekeeping control. One representative experiment is shown (n=4). **C)** RIP-qPCR showing the interaction between hnRNP C and SINV *nsP1* viral RNA. HEK293 WT or vtRNA1 KO clone 2 were infected with SINV at MOI = 0.5 for 48h. Immunoprecipitations were then performed on cell lysates with anti-hnRNP C or mouse IgG1 antibodies. Viral RNA coprecipitating with hnRNP C was quantified by RT-qPCR, and expressed as fraction of input viral RNA levels, and normalized to the IgG isotype control IP (n=3). **D)** RIP-qPCR showing the interaction between ELAVL1 and SINV genomic RNA. RIP-qPCR was performed as in (D) (n=3). Data represents means ± s.d.. Statistical analyses in Figure 5A was performed using a repeated measures one-way ANOVA compared to siCtrl-treated samples on unnormalized, log-transformed data. Testing in Figure 5C-D was performed using a one-sample *t*-test on log-normalized enrichment values compared to WT samples.

To further explore how vtRNAs affect the function of hnRNP C and ELAVL1 as proviral mediators, we tested how the loss of vtRNAs alters the reported ability of hnRNP C and ELAVL1 to interact with viral RNA in the cytosol during SINV infection^48,53–55^. To this end, we immunoprecipitated ELAVL1 and hnRNP C from SINV-infected HEK293 WT or vtRNA1 KO cells and analyzed the abundance of viral transcripts amongst the co-precipitating RNA by RT-qPCR. Both ELAVL1 and hnRNP C strongly interacted with SINV genomic RNA in infected WT cells (Fig. 5C-D, Fig. S5A-B), consistent with previous studies^48,53–55^. In contrast, these interactions were much reduced in vtRNA1 KO cells. This suggests that by facilitating the nucleocytoplasmic redistribution of hnRNP C and ELAVL1, vtRNA1s promote the association of these proteins with viral RNA and favor their proviral activity.

## Discussion

In this study, we have investigated how ‘riboregulation’ by vtRNAs steers the activity of two host proteins to facilitate viral replication. We find that vtRNAs are upregulated upon infection with a wide range of RNA viruses and are required to sustain SINV and EMCV replication. All four vtRNAs interact with the RBPs ELAVL1 and hnRNP C, albeit to different extents, both in homeostatic conditions and in virus-infected cells. Finally, we demonstrate that vtRNAs regulate nuclear-cytoplasmic shuttling of ELAVL1 and hnRNP C, an event that is required for the proviral activity of these proteins. This work thus uncovers a novel molecular mechanism by which vtRNAs promote viral replication.

Previous studies have identified several DNA and RNA viruses that induce vtRNA expression^18,24,38,39^. Our work extends this repertoire to include several viruses that were not previously studied in the context of vtRNA biology, including picornaviruses, alphaviruses, and beta-coronaviruses. We demonstrate that increased vtRNA expression is a late event during infection, requires active viral replication, and is independent of type I IFN signaling. While the precise mechanism of vtRNA transcriptional upregulation remains unknown, some studies suggest that epigenetic rewiring may play a role in vtRNA expression, at least in the context of cancer^60,61^. Whether viral infection can similarly influence the methylation status of vtRNA promoters requires further investigation. Our finding that vtRNAs are proviral factors aligns with earlier studies on EBV, IAV, and AdV infection^24,39,40^. In EBV-infected B cells, vtRNA1-1 (but not vtRNA1-2 and vtRNA1-3) promotes the expression of anti-apoptotic factors, thereby reducing apoptosis and enhancing viral establishment in an MVP-independent manner^24–26^. Similarly, vtRNAs facilitate IAV replication in human cells and in mice by interfering with PKR activation, NF-κB activation, and type I and type III IFN expression^39^. VtRNA2-1 interacts with PKR *in vitro* and in overexpression studies^31–33^. Finally, vtRNA2-1 enhances AdV replication in a PKR-independent manner, namely by promoting nuclear entry of AdV genomes^40^. In stark contrast, vtRNAs limit viral replication during KSHV infection and thus act as an antiviral factor^41,42^. During KSHV-induced host-translational shut-off, decreased expression of the RNA-specific phosphatase DUSP11 causes an accumulation of vtRNAs with their nascent 5’- triphosphate moiety. Triphosphorylated vtRNAs serve as a ligand for RIG-I and activate an antiviral type I IFN response, thereby limiting KSHV replication^41,42^ Although we did not directly investigate the binding of vtRNAs to RIG-I, the loss of vtRNA1-1, vtRNA1-2, and vtRNA1-3 did not have a consistent impact on the type I IFN response during EMCV or SINV infection; if anything the IFN response is increased in infected vtRNA1-deficient cells (Fig. 2M-N). This suggests that vtRNAs are unlikely to play a role in RIG-I activation during SINV and EMCV infection. In addition, during picornavirus infection RIG-I is degraded by viral proteases and therefore cannot sense viral or endogenous RNAs and participate in type I IFN induction in this context^62,63^.

The lack of an overt role for vtRNAs in type I IFN induction or signaling in our experimental set-up is perhaps not surprising. Both SINV and EMCV express strong IFN antagonists that inhibit type I IFN production as well as JAK/STAT signaling downstream of the type I IFN receptor^64,65^. In these contexts, vtRNAs likely affect viral replication more prominently via other cellular pathways. Indeed, blockade of IFN signaling by ruxolitinib did not restore viral replication in vtRNA1-deficient cells to levels seen in WT cells (Fig. 2Q-R). However, we certainly do not exclude that vtRNAs may affect the antiviral IFN response in other viral infections. It would be interesting to use RAP-MS to uncover the vtRNA interactome and potential interactions with ISGs in cells infected with viruses that lack an IFN antagonist. Another possibility is that vtRNAs indirectly regulate type I IFNs through their interactions with cellular proteins. Of note, both hnRNP C and ELAVL1 may modulate the type I IFN response. Loss of hnRNP C-mediated splicing induces the incorporation of *Alu* elements in mRNA, leading to spontaneous activation of the dsRNA sensor MDA5 and subsequent induction of the type I IFN response in breast cancer cells^66^. Additionally, ELAVL1 binds to AU-rich elements in the 3’ UTR of mRNAs coding for IFN-β and multiple ISGs, thereby increasing their stability and translation^67,68^. Thus, the interaction of vtRNAs with these proteins may ultimately lead to downstream effects on the type I IFN response.

Only a few studies have investigated the protein interactome of vtRNAs using pulldowns with *in vitro* transcribed or aptamer-tagged vtRNAs and primarily focused on vtRNA1-1 and vtRNA1-2^25,37^. The advantage of the RAP-MS pipeline is that UV-crosslinking generates a snapshot of endogenous RNA-protein interactions in their natural stoichiometry and allows stringent purification of RNA-protein complexes^46^. The retrieved proteins thus represent high-confidence interactions. Since vtRNAs share a high degree of homology, individual probes may - to some degree - also have interacted with other vtRNA family members. Hence, some of the identified interacting proteins may not be uniquely associating with the target vtRNA, but may rather reflect an interaction with the vtRNA family. Validation of interactions using protein-centric techniques such as RIP-qPCR is therefore crucial. Using RAP-MS and RIP-qPCR, we identified high-confidence interactions between the RBPs hnRNP C and ELAVL1 and vtRNAs in uninfected and infected cells. Both proteins localize to the nucleus and are recruited to the cytoplasm where they facilitate alpha- and picornaviral replication^48,53–55,59^. In the absence of vtRNAs these proteins translocate less efficiently, showing that vtRNAs can regulate protein localization during viral infection. Similarly, a recent study demonstrates that vtRNA1-1 regulates nuclear translocation of the transcription factor TFEB during starvation via the MAPK pathway^26^. Furthermore, vtRNAs promote access of AdV genomes to the nucleus by modulation of the kinesin pathway^40^. Adding to these reports, our observations strengthen the notion that vtRNAs control nucleocytoplasmic transport via multiple mechanisms. How precisely vtRNAs interact with ELAVL1 and hnRNP C to favor their nuclear export remains to be determined, for example through identification of the exact interaction interface or by live cell imaging. In addition, it is unclear why vtRNAs interact with hnRNP C and ELAVL1 under homeostatic conditions without altering their localization. We speculate that viral proteins initiate changes in the nucleocytoplasmic distribution of proteins and that the increased expression of vtRNAs during viral infection is needed to put the balance in favor of cytosolic translocation. Interestingly, we identified several other proteins (hnRNP M, XRCC5, and PTBP1, Fig. 3B) in the vtRNA interactome that promote SINV replication^53–55^. These proteins localize to the nucleus in uninfected cells yet interact with SINV RNA, implying they are shuttled to the cytosol, analogous to hnRNP C and ELAVL1^53–55^. It is therefore an interesting possibility that vtRNAs may similarly contribute to the cellular redistribution of these proteins to enhance viral infection.

ELAVL1 was recently reported to interact with vtRNA2-1^30^. The expression of vtRNA2-1/mvtRNA is upregulated in the colon or ileal mucosa of patients with ulcerative colitis or Crohn’s disease or in mice treated with colitis-inducing agents^30^. By binding ELAVL1, vtRNA2-1 prevents the stable association of ELAVL1 with several mRNAs encoding proteins that mediate epithelial barrier integrity, leading to decreased barrier function. We found that ELAVL1 not only interacts with vtRNA2-1, but even more strongly with vtRNA1-1, vtRNA1-2, and vtRNA1-3, which persists during viral infection.

Since loss of epithelial barrier integrity favors access of a virus to the underlying tissue, it is interesting to speculate whether the virus-induced upregulation of vtRNAs and their interaction with ELAVL1 may promote viral spread by impairing barrier function, consistent with the proviral activity of these interaction partners. This could be addressed in organotypic model systems or *in vivo* in a recently described mvtRNA-deficient mouse model^69^.

VtRNAs derive their name from their association with the enigmatic vault complex. An estimated 5% of cellular vtRNAs are associated with the vault complex^13,18,31^. MVP is not located in the nucleus, and kinetics of nucleocytoplasmic trafficking of proteins with a classical nuclear localization signal or nuclear export signal are unaffected in MVP-deficient mouse embryonic fibroblasts^70^. This suggests that the subcellular localization of ELAVL1 and hnRNP C may be regulated by ‘free’ vtRNAs. Nevertheless, vtRNAs may exert other proviral functions in association with the vault complex, which remains to be established.

The strong upregulation of vtRNAs during various viral infections and their proviral function in multiple infection models make these ncRNAs potential druggable targets to treat infectious diseases. Our study adds insight into the molecular mechanisms of vtRNAs during such diseases and thus brings this goal closer within reach.

## Acknowledgements

We are grateful to Sebenzile Myeni and Marissa Linger for help with MERS-CoV infections, Linda Boomaars-van der Zanden for help with MHV infections, and Diana van den Wollenberg and Rob Hoeben for help with adenovirus and reovirus infections. We thank Lennard Voortman for advice on microscopy experiments. We thank all members of the Van der Veen group as well as Fiamma Salerno and Maaike Ressing for assistance and critical discussions. This work was supported by a research grant from the Institute for Chemical Immunology (ICI-00203), which is funded by a Gravitation project from the Netherlands Organization for Scientific Research (NWO), a Vidi research grant and an Aspasia award from the NWO (09150171910070), an ENW-M1 grant from the NWO, and the LUMC.

---

**Supplemental Figure 1, belonging to Figure 1.**
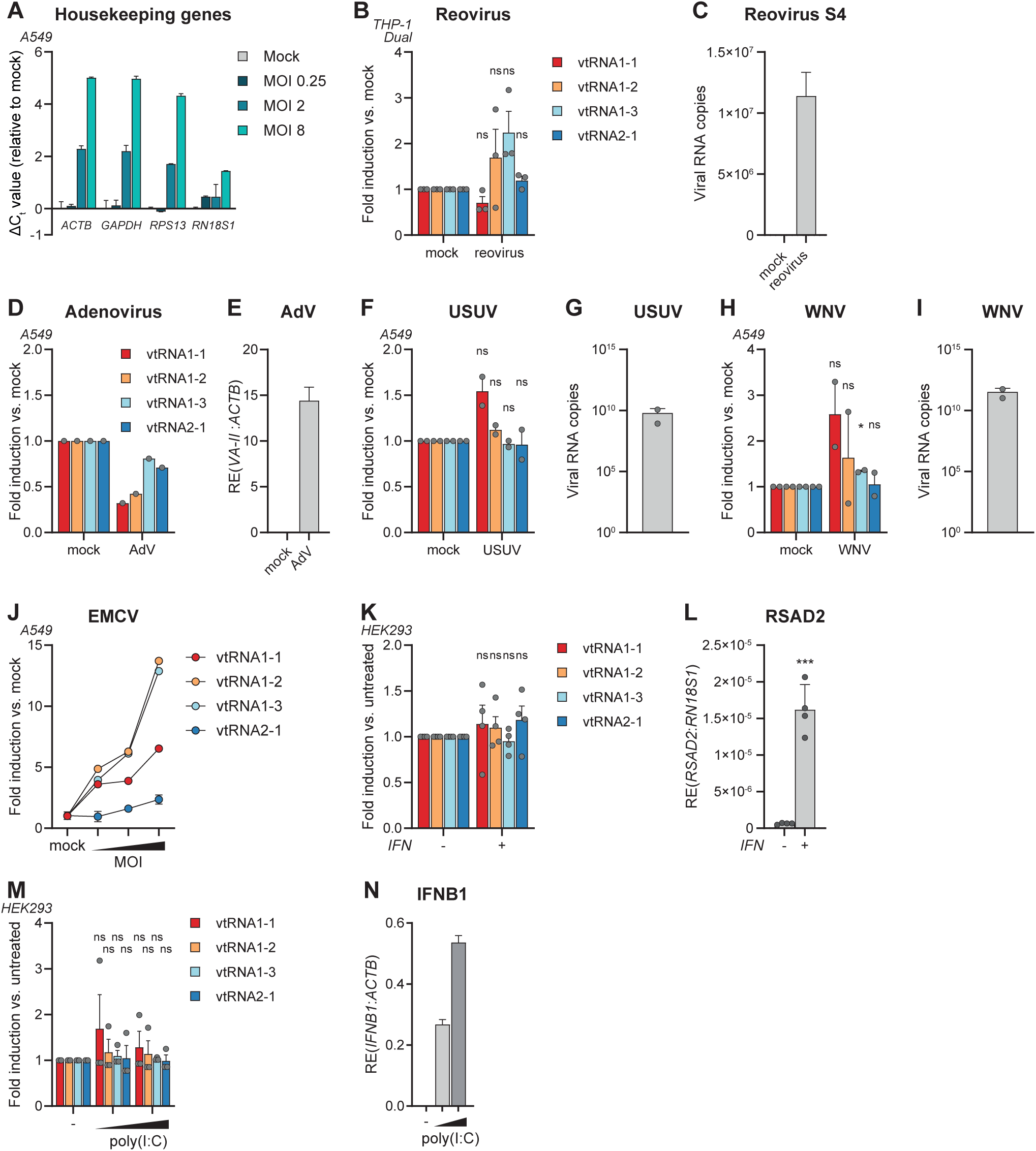
**A)** Expression of common housekeeping genes were measured in A549 cells 24hpi with increasing doses of EMCV by RT-qPCR. The difference in C_t_ value (ΔC_t_) compared to mock-infected cells is plotted. One representative graph is shown (n=2). **B)** THP-1 Dual monocytes were PMA-differentiated into macrophages and infected with reovirus (MOI = 10, 24hpi). Expression of vtRNA1-1, vtRNA1-2, vtRNA1-3 and vtRNA2-1 was measured using RT-qPCR and normalized against expression of *ACTB*. Values are displayed relative to mock-infected cells (n=3). **C)** The amount of reovirus *S4* copies per well in (B) was determined by RT-qPCR. One representative image is shown (n=3). **D)** A549 cells were infected with human adenovirus (AdV; MOI = 10, 24hpi). Expression of vtRNAs was measured as in (B). Values are displayed relative to mock-infected cells. **E)** Expression of the *VA-II* RNA of human AdV was measured by RT-qPCR to show productive infection in (D). Relative expression was normalized against *ACTB*. **F)** A549 cells were infected with Usutu virus (USUV; MOI = 3, 24hpi). Expression of vtRNAs was measured by RT-qPCR and normalized against expression of *RN18S1*. Values are displayed relative to mock-infected cells (n=2). **G)** The amount of intracellular USUV copies was determined by RT-qPCR to show productive infection (n=2). **H)** A549 cells were infected with West Nile virus (WNV; MOI = 1, 24hpi). Expression of vtRNAs was measured as in (F). Values are displayed relative to mock-infected cells (n=2). **I)** The amount of intracellular WNV copies was determined by RT-qPCR to show productive infection (n=2). **J)** A549 cells were infected with increasing doses of EMCV (MOI = 0.25, 2, and 8). Expression of vtRNAs was measured as in (F). One representative graph is shown (n=3). **K)** HEK293 cells were stimulated with 200 U/ml recombinant type I IFN for 24h. Expression of vtRNAs was measured as in (F). Values are displayed relative to untreated cells (n=4). **L)** Expression of the interferon-stimulated gene *RSAD2* in untreated versus IFN-treated cells in (K) was measured by RT-qPCR (n=4). **M)** HEK293 cells were transfected with increasing doses of the viral dsRNA mimic poly(I:C) (75 or 125 ng). After 24h, cells were harvested and expression of vtRNAs was measured as in (F). Values are displayed relative to untreated cells (n=3). **N)** Expression of IFN-β mRNA following poly(I:C) transfection in (M) was analyzed by RT-qPCR. Relative expression was calculated as in (B). One representative graph is shown (n=3). Data represents means ± s.d.. Statistical analysis in S1B, S1F, S1H, and S1K was performed on unnormalized, log-transformed data using a Student’s *t*-test to test significance of each vtRNA expression between virus-infected/treated and mock-infected/untreated conditions. In S1M, a repeated measures one-way ANOVA was performed.

**Supplemental Figure 2, belonging to Figure 2.**
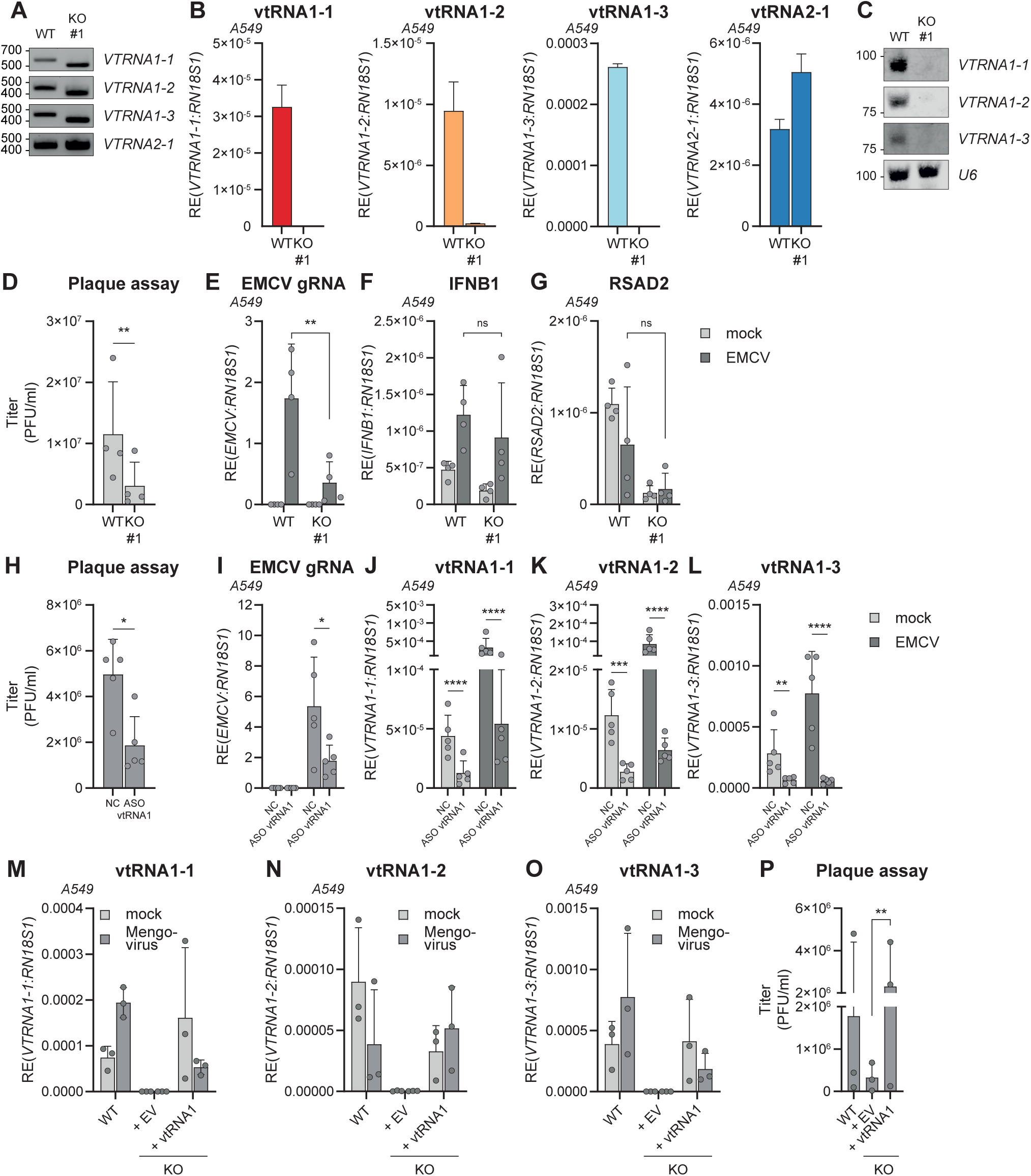
**A)** Primers flanking each *VTRNA* gene were designed to assess the integrity of the gene after CRISPR-Cas9 genome editing, compared to A549 WT cells. The monoclonal KO cell line (#1) was chosen based on disruption of the *VTRNA1* loci. One representative experiment is shown (n=2). **B)** Loss of vtRNA expression in the A549 KO clone selected in (A) was shown using RT-qPCR. Expression of vtRNAs was measured and normalized to *RN18S1*. Data represent means ± s.d.; one representative experiment is shown (n=4). **C)** Northern Blot showing the loss of vtRNA1-1, vtRNA1-2, and vtRNA1-3 expression in the vtRNA1 KO clone selected in (B). U6 snRNA expression serves as housekeeping control. **D)** A549 WT or vtRNA1 KO cells were infected with EMCV for 16h. Viral titer was determined by plaque assay (n=4). **E)** RT-qPCR showing the relative expression of EMCV genomic RNA (gRNA) in cells infected in (D), measured with primers targeting the EMCV *Leader* gene. Data are normalized against *RN18S1* (n=4). **F)** RT-qPCR showing the relative expression of *IFNB1* in cells infected in (D), normalized against *RN18S1* (n=4). **G)** RT-qPCR showing the relative expression of *RSAD2* in cells infected in (D), normalized against *RN18S1* (n=4). **H)** A549 WT cells were treated with ASOs against vtRNA1-1, vtRNA1-2, and vtRNA1-3 (‘ASO vtRNA1’) or a non-targeting control (‘NC’) for 48h. Cells were subsequently infected with EMCV for 20h, and viral titers were determined by plaque assay (n=5). **I)** RT-qPCR showing the relative expression of EMCV gRNA in cells infected in (H), normalized against *RN18S1* (n=5). **J-L)** RT-qPCRs showing the relative expression of vtRNA1-1, vtRNA1-2, and vtRNA1-3 following ASO-mediated knockdown in A549 cells and subsequent EMCV infection (n=5). **M-O)** RT-qPCRs showing the relative expression of vtRNA1-1, vtRNA1-2, and vtRNA1-3 in A549 WT or vtRNA1 KO cells. VtRNA1 KO cells were transduced with empty lentiviruses (EV) or lentiviruses encoding *VTRNA1-1*, *VTRNA1-2* and *VTRNA1-3*. Cells were mock-infected or infected with Mengovirus for 20h. Relative expression was normalized against *RN18S1* (n=3). **P)** Mengovirus titers in cells treated as described in (M-O) were determined using a plaque assay (n=3). Data represents means ± s.d. Statistical analyses in S2D-I and S2P were performed on unnormalized, log-transformed data using a Student’s *t*-test. Data in S2J-L were analyzed using a repeated measures one-way ANOVA.

**Supplemental Figure 3, belonging to Figure 3.**
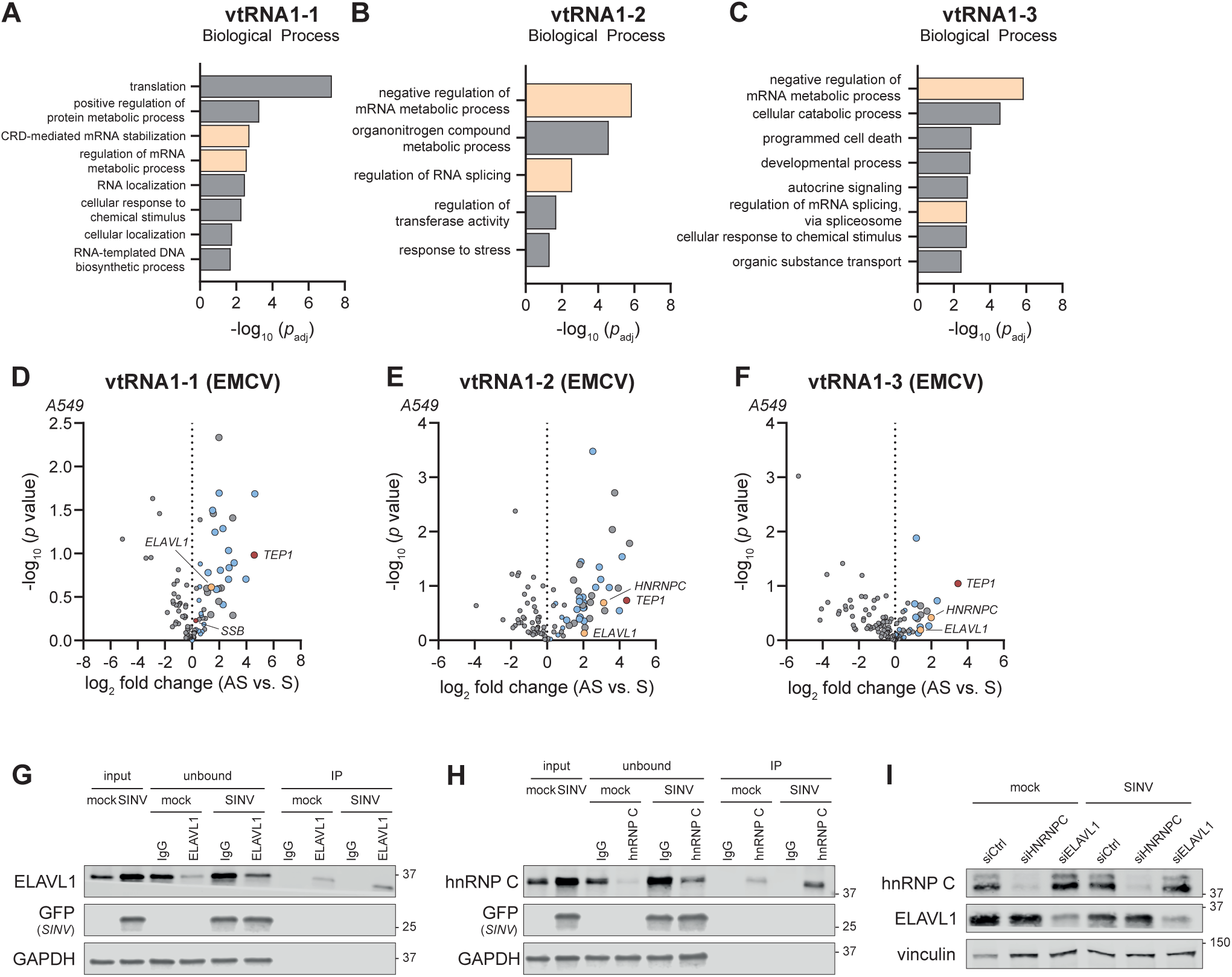
**A-C)** GO Term analysis showing the enriched Biological Processes of interactors of vtRNA1-1, vtRNA1-2, and vtRNA1-3 in uninfected A549 cells. **D-F)** Quantification of vtRNA-interacting proteins (antisense, ‘AS’) compared to the negative control (sense, ‘S’) in cells infected with EMCV (MOI = 0.25, 24 hpi), expressed as log_2_-fold change. The enriched proteins that are classified as “RNA-binding” according to GO Term analysis are highlighted in blue. Positive controls are highlighted in red, and proteins of interest are highlighted in yellow. Statistical analysis was performed using a two-tailed *t*-test with a *post-hoc* FDR correction (n=2). **G-H)** The efficiency of immunoprecipitation of ELAVL1 and hnRNP C, described in Fig. 3M-N, was demonstrated by immunoblotting. GFP staining indicates samples that were infected with SINV. GAPDH staining serves as a housekeeping marker to show equal loading in the IgG and ELAVL1 IP. Representative image is shown (n=3). **I)** The efficiency of siRNA-mediated knockdown of hnRNP C and ELAVL1 in Fig. 3O was determined by immunoblotting. Vinculin serves as a housekeeping marker. Representative image is shown; data represents means ± s.d. (n=2).

**Supplemental Figure 4, belonging to Figure 4.**
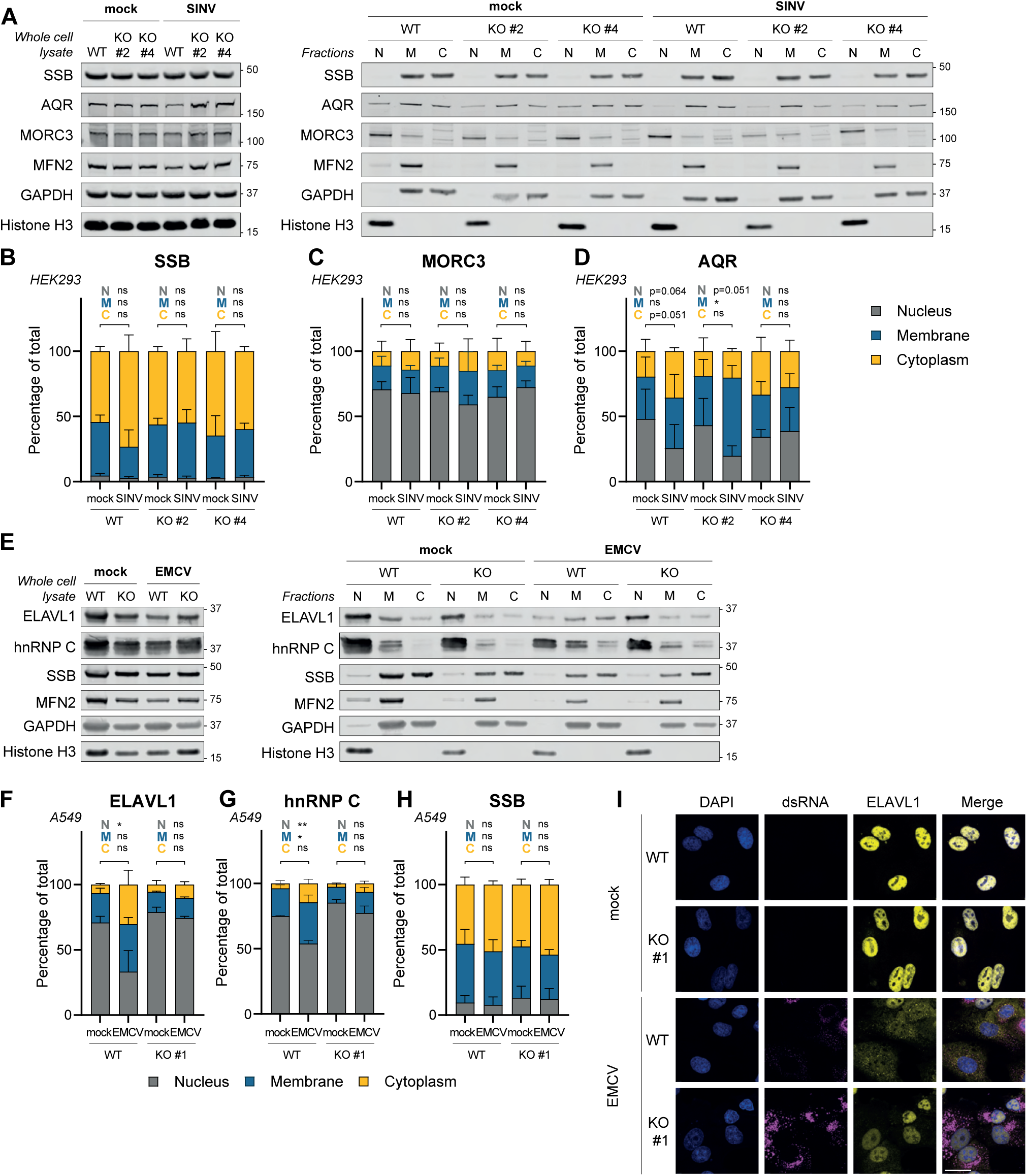
**A)** The expression of SSB, AQR and MORC3 in whole cell lysate (left) or different subcellular compartments (right). Cell lysates of uninfected or SINV-infected HEK293 WT or vtRNA1 KO cells (clone 2 and 4) were fractionated into a nuclear (N), membranous (M) or cytoplasmic (C) fraction and analyzed by immunoblotting. of uninfected or SINV-infected HEK293 WT or vtRNA1 KO cells (clone 2 and 4). GAPDH, mitofusin 2 (MFN2), and histone H3 are used as markers for the cytoplasmic/membranous, membranous, and nuclear fraction, respectively. One representative experiment is shown (n=3). **B-D)** Quantification of SSB, MORC3 and AQR band intensities in (A). Intensity of each band was expressed as a percentage of cumulative intensity of the three fractions within each condition (n=3); *p* values for statistical tests between every fraction are shown preceding with ‘N’ (nucleus), ‘M’ (membrane), or ‘C’ (cytoplasm). **E)** Expression of ELAVL1, hnRNP C and SSB in whole cell lysate (left) or different subcellular compartments (right). Cell lysates of uninfected or EMCV-infected A549 WT or vtRNA1 KO cells were fractionated into a nuclear (N), membranous (M) or cytoplasmic (C) fraction and analyzed by immunoblotting. GAPDH, mitofusin 2 (MFN2), and histone H3 are used as markers for the cytoplasmic/membranous, membranous, and nuclear fraction, respectively. One representative experiment is shown (n=2). **F-H)** Quantification of ELAVL1, hnRNP C, and SSB band intensities in (E). Intensity of each band was expressed as a percentage of cumulative intensity of the three fractions within each condition (n=2); *p* values for statistical tests between every fraction are shown preceding with ‘N’ (nucleus), ‘M’ (membrane), or ‘C’ (cytoplasm). **I)** A549 WT or vtRNA1 KO cells were fixed and permeabilized 24h after infection with EMCV, stained for ELAVL1 (yellow) and dsRNA (magenta) and analyzed by immunofluorescence microscopy. Nuclei were stained with DAPI (blue). Scale bar indicates 25 µm. Representative micrographs are shown (n=2). Data in stacked bar graphs represent means ± s.d. Statistical analysis in S4B-D and S4F-H were performed using a repeated measures one-way ANOVA on unnormalized data.

**Supplemental Figure 5, belonging to Figure 5.**
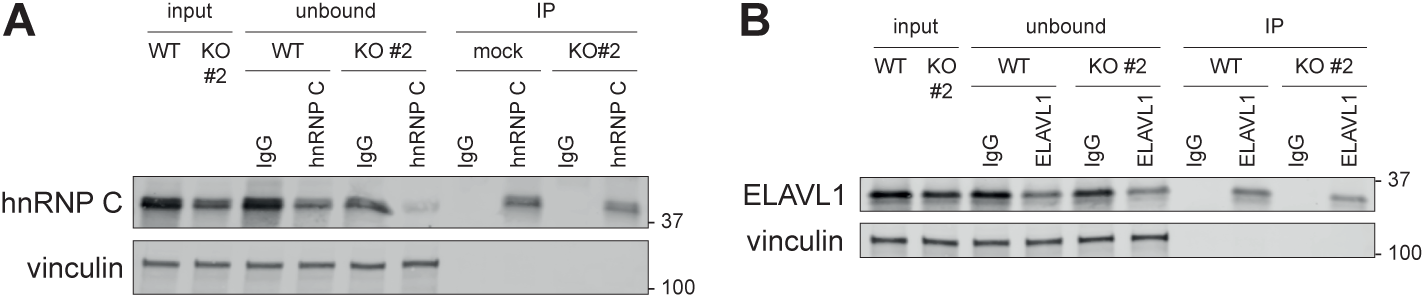
**A-B)** The efficiency of immunoprecipitation of hnRNP C and ELAVL1, described in Fig. 5C-D, was determined by immunoblotting. Vinculin staining serves as a housekeeping marker to show equal loading in the IgG and ELAVL1 IP. Representative image is shown (n=3).

